# Wheat shovelomics I: A field phenotyping approach for characterising the structure and function of root systems in tillering species

**DOI:** 10.1101/280875

**Authors:** Larry M. York, Shaunagh Slack, Malcolm J Bennett, M John Foulkes

**Keywords:** Rooting, abiotic stress, Triticum aestivum L, fertilizer, water, nitrogen, soil, resource acquisition

## Abstract

Wheat represents a major crop, yet the current rate of yield improvement is insufficient to meet its projected global food demand. Breeding root systems more efficient for water and nitrogen capture represents a promising avenue for accelerating yield gains. Root crown phenotyping, or shovelomics, relies on excavation of the upper portions of root systems in the field and measuring root properties such as numbers, angles, densities and lengths. We report a new shovelomics method that images the whole wheat root crown, then partitions it into the main shoot and tillers for more intensive phenotyping. Root crowns were phenotyped using the new method from the Rialto × Savannah population consisting of both parents and 94 doubled-haploid lines. For the whole root crown, the main shoot, and tillers, root phenes including nodal root number, growth angle, length, and diameter were measured. Substantial variation and heritability were observed for all phenes. Principal component analysis revealed latent constructs that imply pleiotropic genetic control of several related root phenes. Correlational analysis revealed that nodal root number and growth angle correlate among the whole crown, main shoot, and tillers, indicating shared genetic control among those organs. We conclude that this phenomics approach will be useful for breeding ideotype root systems in tillering species.

## Introduction

The global population is expected to increase to nine billion people by 2050 which necessitates an increase in global food production of at least 60%, but likely as much as 100% due to increased livestock production (Grafton *et al.*, 2015). Wheat is a staple crop in many countries, grown on 220 million hectares with a global yield of 729 million tonnes (FAOSTAT, 2014). However, the current yield improvement rate of 1% for wheat is not sufficient to meet greater demands of the future global population (Ray *et al.*, 2013). Planting more hectares is not a viable option (Pretty, 2008). Similarly, neither is adding more fertilizer due to water and atmospheric pollution (Jenkinson, 2001). In order to increase wheat yields while mitigating environmental degradation, varieties are needed with greater water and nitrogen uptake capacity.

Worldwide, drought limits wheat yields more than any other single factor and the ability to acquire water is principally related to the size of the plant’s root system (Cattevelli *et al*., 2008; Wasson *et al*, 2012). Nitrogen acquisition efficiency is an important component of crop nitrogen use, and is generally defined as the ability of the plant root system to uptake nitrogen from the soil. The wheat root system begins with the primary root and seminal roots that arise from the scutellar and epiblast nodes (Klepper *et al.*, 1984), which are referred to as the tap and basal roots in the terminology proposed by the International Society for Root Research (ISSR; Zobel and Waisel, 2010). As wheat develops, both roots and tillers emerge from the coleoptile node and subsequently the leaf nodes. These shoot-borne roots, as they are termed by ISRR, are widely referred to as crown or nodal roots. These nodes contain four positions from which either a root or a tiller can emerge. The main shoot can produce several nodes with tillers and roots, and tillers can produce their own nodes with sub-tillers and roots. Through this developmental process, wheat produces a complex axial root system of tillers and crown roots (Klepper *et al.*, 1984). The mature wheat root system can spread 30 – 90 cm laterally from the stem (Manschadi *et al.*, 2006) and grow to a maximum depth of 1 – 2 m (Kirkegaard and Lilley, 2007). The root crown refers to the entirety of only the excavated root system in the surface soil layer (ca. 0 −20 cm soil depth). Relatively little is known about how genotypic variation in these surface roots relates to variation in the root system below the root crown, or how this variation could affect soil resource acquisition.

Root system architecture (RSA) refers to the explicit three-dimensional spatial configuration of all root axes in a root system (Lynch, 1995). Root phenomics is an emerging field that leverages high-throughput phenotyping to measure properties of hundreds of individual root systems. An elemental unit of phenotype measured is referred to as a phene (Lynch and Brown, 2012; Serebrovsky, 1925), with the analogy ‘phene is to phenotype as gene is to genotype.’ The word phene completely replaces the ambiguously used term ‘trait’ (Violle *et al.*, 2007). RSA phenes include numbers, lengths, angles, and diameters of different classes of roots (Lynch, 1995). The term phene state is used to denote a specific value of a measured phene (York *et al.*, 2013), for example the phene state for crown root number could be 40. Phene aggregates refer to other phenotypic measures that combine the values of more elemental phenes. For example, root length density is a phene aggregate that combines the numbers and lengths of many classes of roots (York *et al.*, 2013). A phene-based paradigm is needed to fully understand root phenotypes and their relation to crop performance (York *et al.*, 2013).

An optimal root length density of 1 cm cm^-3^ is predicted for wheat uptake of water and nitrate (Foulkes *et al.*, 2009; van Noordwijk, 1983), so RSA phenes that possibly influence root length density such as shoot number, nodal root number, and lateral root branching density are hypothesized to be important for soil resource acquisition. Research on maize (*Zea mays*) likewise suggests phenes that optimize root density such as nodal root number (Saengwilai *et al.*, 2014; York *et al.*, 2013), lateral branching density (Postma *et al.*, 2014; Zhan and Lynch, 2015), and the number of root nodes (York and Lynch, 2015) are influential for water and nitrate uptake. Angles of nodal roots have also been associated with both deep and shallow root foraging (Dathe *et al.*, 2016; Trachsel *et al.*, 2013; York and Lynch, 2015). High-throughput phenotyping of wheat RSA in the field is critical for making associations among root phenes, genes, and functional utility and for the development of molecular markers for deployment in plant breeding.

Phenotyping wheat root system architecture is advancing, but is often limited to seedling screens in clear pots (Richard *et al.*, 2015) or on germination paper (Atkinson *et al.*, 2015; Bai *et al.*, 2013). While the seminal root system is undoubtedly important for early establishment, the post-embryonic root system dominates later in growth and little evidence exists for whether or not mature root system properties can be predicted from seedling properties. Soil coring is the most common method used in the field to quantify the distribution of root length with depth nearer to crop maturity, either applying the soil-core break method (Wasson *et al.*, 2014) or root washing and image analysis (Ford *et al.*, 2006; White *et al.*, 2015). However, no information is gained about the underlying root system architecture that determines root density with depth. Recently, high-throughput phenotyping of wheat RSA was accomplished in the field in durum wheat (Maccaferri *et al.*, 2016) using a modified ‘shovelomics’ method developed in maize (Trachsel *et al.*, 2011) where entire wheat root crowns were excavated, washed, and imaged before being processed with REST automatic image processing software (Colombi *et al.*, 2015). The shovelomics approach is promising. However, the previous method did not analyse the influence of tillering on wheat root system architecture, nor were the number of nodal roots counted. Whether the main shoot roots or tiller roots influence the whole root crown equally is not known, nor is it known whether the phenes correlate among the tillers. Given the importance of tillering for wheat production, a more complete account of wheat root system architecture that quantifies RSA among tillers is required. Here, a new phenomics approach is described for intensive phenotyping of wheat root system architecture, investigating relations among tiller and main shoot rooting phenes, and calculating heritabilities for root phenes that are important for soil resource acquisition. We hypothesized root phenes would correlate among main shoot root systems and tiller root systems and that measurements of main shoot root system phenes would provide greater heritability than whole crown phenes.

## Methods

### Plant material

The germplasm consisted of two winter wheat parents, Rialto and Savannah, and 94 doubled-haploid lines developed from the F_1_ cross (Atkinson *et al.*, 2015). Both parents are semi-dwarf (*Rht-D1b*, formerly *Rht-2*) UK winter wheat, and hard endosperm cultivars. Rialto is suitable for some bread-making processes, and was bred by RAGT Ltd and released in 1995. Savannah is a feed wheat cultivar bred by Limagrain UK Ltd with high yield potential, and released in 1998.

### Field experiments

The field site was at the University of Nottingham farm in Sutton Bonington, United Kingdom (52°50 N, 1°14 W). The experimental design was a randomized complete block design with genotypes randomized within blocks and four replicates. Plots were 6 × 1.65 m and planted with 320 seeds m^-2^ for a target spring density of 200 plants m^-2^. The experiment was planted on October 20, 2014. The soil was a sandy medium loam to 80 cm over Kyper marl clay of the Dunnington Heath series. Two blocks were irrigated using drip tape, and the other two were rainfed. For the current analyses, the water treatment was ignored as the changes in root crown phene states were not substantial because of adequate rainfall.

### Wheat shovelomics

Root crowns were excavated approximately two weeks after anthesis on July 8-9, 2015. Several (four to eight) adjacent plants per plot were selected from an internal row based on their uniform size and presence of neighbouring plants. A straight-edged spade with a width of 15 cm was inserted directly adjacent to the neighbouring rows on either side of the focal plants with the width of the blade parallel to the row. Insertion was completely perpendicular to the ground. The focal plants and attached soil were lifted from the ground on the spade and placed into a plastic bag taking care to not let the heavy soil tear off the fragile roots. Sample bags with entire excavated plants were transported to a nearby washing station where roots with attached soil were carefully lifted out of the bag and placed into 10 litre buckets filled with water. The samples were left soaking in the water at least 10 minutes to allow the soil to loosen. Following this, root crowns were gently moved back and forth in the water to facilitate soil removal, then lifted out and sprayed with low pressure water from a hose to remove the rest of the soil. The root crowns of each individual plant from the group excavated form the plot were evaluated visually, then a single root crown of average overall size and average root diameters was retained for subsequent analysis, while the others were discarded. The root crowns were severed from the shoots close to the base, then root crowns (with ca. 1 cm of the basal shoots attached; Figure. 1) were placed back in the bag for cold storage at 5 ° C.

**Fig. 1.**
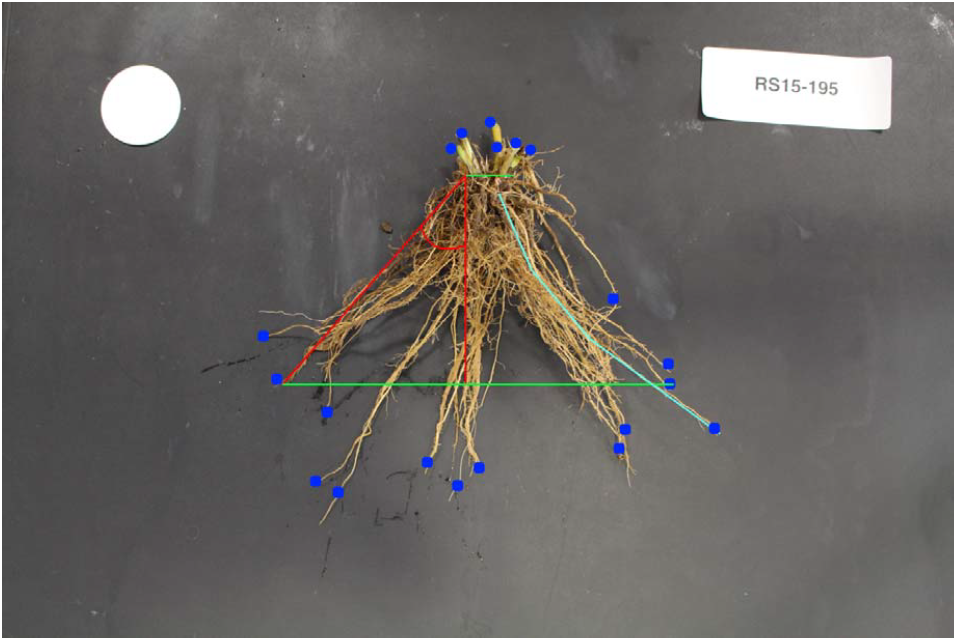
An example image of the entire wheat root crown (centre), circular scale (top left), and plot ID tag (top right). Some of the measured features include stem width (top green line), system width (bottom green line), the nodal root growth angle derived trigonometrically from stem and system widths (red arc), number of tillers (upper blue points), number of nodal roots (lower blue points), and nodal root length (light blue polyline).

Root crowns were imaged using a Canon EOS 70D DSLR camera attached to an aluminium frame covered in black cloth but with an open front in order to minimize directional lighting and maximize diffuse lighting in otherwise ambient lighting of a laboratory. The camera was a consumer Canon camera with manual settings for the shutter time duration and aperture to optimize the root contrast with background. The root crowns were placed on a matte black vinyl background along with a 42 mm circular filter paper and a white tag printed with the sample ID. The whole root crown was imaged first, then split into the main shoot and tillers. The main shoot was identified by presence of the seminal root system and imaged next, followed by all tillers sorted by apparent size which is presumed to be indicative of the tiller appearance order. A column was added to the resulting data set that included the plot number, then results were compiled at the whole crown level based on aggregating within the plot.

Images were analysed using a modified method from York and Lynch (2015). A project for the *ObjectJ* plugin (https://sils.fnwi.uva.nl/bcb/objectj) for *ImageJ* (Schneider *et al.*, 2012) was created to allow the angles, numbers, and lengths of nodal roots to be measured from the whole crown, main shoot, and all tillers (Figure 1). The pixel dimensions were converted to physical units using measurements of the known-sized scale in every image. Table 1 defines all measured phenes. A polyline was used to measure the crown lengths of the outermost roots and the seminal root length, and the angle was measured for the outermost crown roots at approximately 5 cm depth by measuring the width then later calculating angle using trigonometry and the actual depth measurement to where width was measured. For nodal root number, each nodal root axis was manually annotated and the count recorded in an output file. The image analysis gave values for the number of pixels corresponding to root length and numbers. Using the 42 mm circular scale, these pixel values were then converted to the relevant units for each root measurement using a script written in *R* (R Core Team, 2012) that also calculated angles as detailed in York *et al.* (2015).

**Table 1.** A list of traits measured in this study, which section of a crown measured from, and their derivations. Measurements of only the whole crown are aggregated from the individual shoots and tillers (Whole), measurements done on all shoots and the crown (All), and measurements only done on the individual shoots (shoots).

| Trait | Section | Definition |
| --- | --- | --- |
| Tiller number | Whole | Aggregated from the number of shoot images for each crown |
| Total nodal root number | Whole | Aggregated from counts on individual shoots |
| Stem width | All | On crown, width at base of all stems, on shoots width of stem |
| System width | All | Width of the root system at widest point with roots present on both sides |
| Depth-to-width | All | The vertical distance from stem width to system width |
| Nodal root growth angle | All | Derived from stem width, system width, and depth-to-width |
| Nodal root length | All | Length of polyline tracing longest crown roots on left and right |
| Nodal root diameter | Shoots | Width at widest point of representative nodal root |

### Statistics

Statistics were performed using R version 3.3.2. Root crowns were identified as those of the whole crown, main shoot, and tillers based on their image order while having the same plot identification (ie, the whole root crown was separated into component root crowns of the different shoots). Based on the same principle, the total number of nodal roots for the whole crown and the shoot numberwas calculated by aggregating data within a single root and shoot sample. Broad-sense heritability was calculated based on (Falconer and Mackay, 1996) as:

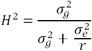

The variables 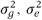 and r represent the variance of the genotype effect, variance of the environment effect, and the number of replicates (here, 4), respectively. The variances were obtained by fitting a mixed model including genotype as a random effect and replicate as a fixed effect using the *lme4* package.

For principal component analyses, data were centered and scaled using the *scale* function, and then the *prcomp* function was used to conduct the principal component analysis (PCA). Cronbach’s α was used as a measure of internal consistency and was calculated using the *alpha* function from the *psych* package.

## Results

### Heritable genotypic variation

Substantial variation existed for all measured phenes (Figure 2, Table 2). Differences were observed among population averages for the whole crown, main shoot, and tillers for several phenes, especially those related to root crown size, such as root system width and depth to width ratio. Heritabilities ranged between 0 - 0.52 and were generally greater for the whole crown phenes (Table 2). Correlations among root phenes were common, and especially strong when correlating the same phenes of tillers to the main shoot (Figure 3).

**Table 2.** Traits from the whole crown (cr), main shoot (ms), and tillers 1-3 (t1-t3) with means of all samples, minimums, maximums, F-values from ANOVA of genotype effect, significance level, and broad-sense heritability. Significance levels displayed as ns >. 05, *<.05 >.01, ** <.01.

| Trait | Mean | Minimum | Maximum | F-value | H<sup>2</sup> |
| --- | --- | --- | --- | --- | --- |
| cr. stem width mm | 19.18 | 3.27 | 60.28 | 1.38 * | 0.29 |
| cr. system width mm | 67.93 | 28.17 | 196.00 | 1.61 ** | 0.35 |
| cr. depth to width mm | 38.43 | 15.50 | 101.06 | 1.25 ns | 0.17 |
| cr. crown length mm | 110.49 | 58.37 | 215.74 | 1.16 ns | 0.14 |
| cr. angle ° | 57.51 | 28.01 | 82.06 | 1.58 ** | 0.37 |
| cr. total NRN | 38.82 | 12.00 | 142.00 | 2.02 ** | 0.51 |
| cr. tiller num | 5.52 | 2.00 | 24.00 | 2.04 ** | 0.52 |
| cr. seminal length mm | 180.45 | 89.52 | 327.93 | 1.02 ns | 0.02 |
| ms. stem width mm | 4.98 | 2.04 | 12.59 | 1.06 ns | 0.07 |
| ms. system width mm | 53.00 | 17.15 | 151.50 | 1.44 * | 0.26 |
| ms. depth tow idth mm | 36.96 | 7.88 | 94.65 | 1.17 ns | 0.10 |
| ms. crownlength mm | 101.09 | 45.17 | 187.98 | 1.04 ns | 0.06 |
| ms. angle ° | 56.19 | 16.78 | 78.96 | 1.05 ns | 0.07 |
| ms. nodal root number | 9.13 | 4.00 | 16.00 | 1.38 * | 0.28 |
| ms. nodal root diam mm | 1.57 | 0.35 | 3.48 | 1.25 ns | 0.20 |
| t1. stem width mm | 4.75 | 1.77 | 10.23 | 1.11 ns | 0.09 |
| t1. system width mm | 48.27 | 14.95 | 110.14 | 1.31 * | 0.24 |
| t1. depth to width mm | 31.78 | 9.54 | 98.60 | 1.28 ns | 0.19 |
| t1. crown length mm | 92.35 | 35.15 | 183.29 | 1 ns | 0.00 |
| t1. angle ° | 55.36 | 14.38 | 79.24 | 1.12 ns | 0.11 |
| t1.nodal root number | 8.01 | 4.00 | 17.00 | 1.55 ** | 0.35 |
| t1.nodal root diam mm | 1.54 | 0.39 | 3.64 | 0.83 ns | 0.00 |
| t2. stem width mm | 4.66 | 2.42 | 17.44 | 1.03 ns | 0.00 |
| t2. system width mm | 44.69 | 5.93 | 134.64 | 1.3 ns | 0.24 |
| t2. depth to width mm | 30.31 | 10.19 | 68.44 | 1.12 ns | 0.06 |
| t2. crown length mm | 89.02 | 29.35 | 160.60 | 0.92 ns | 0.00 |
| t2. angle ° | 56.55 | 27.05 | 86.70 | 0.95 ns | 0.00 |
| t2. nodal root number | 7.02 | 2.00 | 13.00 | 1.12 ns | 0.15 |
| t2. nodal root diam mm | 1.53 | 0.47 | 4.03 | 1.32 * | 0.26 |
| t3. stem width mm | 4.42 | 1.97 | 8.24 | 1.14 ns | 0.06 |
| t3. systemwidth mm | 43.01 | 2.66 | 153.27 | 2.06 ** | 0.48 |
| t3. depth to width mm | 27.21 | 2.88 | 95.27 | 1.37 * | 0.00 |
| t3. crown length mm | 86.26 | 25.20 | 172.96 | 1.39 * | 0.32 |
| t3. angle ° | 54.97 | 20.05 | 92.25 | 1.24 ns | 0.25 |
| t3. nodal root number | 6.55 | 2.00 | 14.00 | 1.32 ns | 0.31 |
| t3. nodal root diam mm | 1.52 | 0.34 | 3.51 | 0.8 ns | 0.00 |
| PC1 | 0.00 | -4.65 | 7.49 | 1.43 * | 0.29 |
| PC2 | 0.00 | -3.73 | 4.07 | 1.24 ns | 0.19 |

**Fig. 2.**
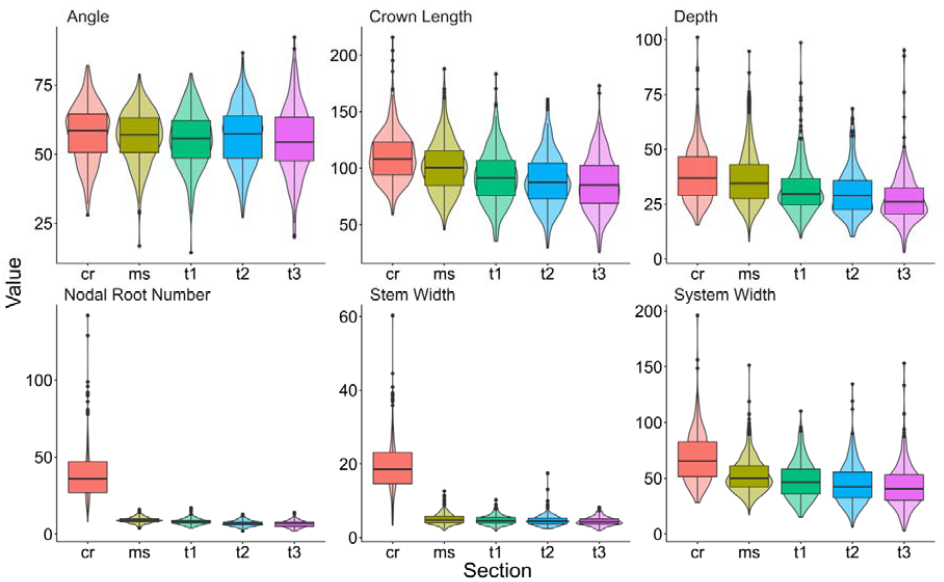
Variation of traits (nodal root angle°, crown root length mm, root system depth mm, nodal root number, stem width mm and root system width mm) and as measured on the entire root crown (cr), main shoot (ms), and tillers 1-3 (t1-t3) from unaveraged data. Box plots with mean and quantiles are overlaid on violin plots giving the frequency distribution.

**Fig. 3.**
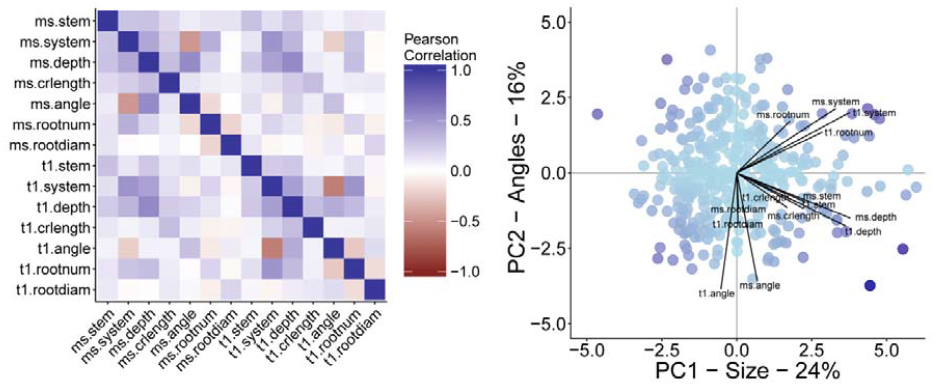
Pairwise correlations of main shoot and first tiller are depicted as a heat map with darker blue indicating greater positive correlations, and darker red indicating greater negative correlations. Principal component analysis (PCA) was conducted for the same 14 traits. A biplot from the PCA depicts a scatterplot of the scores for principal components 1 and 2 (PC1 and PC2) for individual root crowns. The length of labeled lines along an axis indicates the magnitude of correlation between original values of a trait and the principal component. Maximum correlation is 0.77.

### Root number phenes for whole crown, main shoot and tillers

Nodal root number of the whole crown ranged from 12 to 142 with an average of 38.8. The main shoot had an average of 9 nodal roots with tillers 1, 2, and 3 having an average of 8.0, 7.0, and 6.5 nodal roots, respectively. The Pearson correlation was 0.19 (p <. 001). between nodal root numbers on the main shoot versus tiller 1, while the average inter-item correlation of angles of the whole crown, main shoot, and tillers 1-3 was 0.28. Cronbach’s α was used as a measure of internal consistency of all these root numbers and was determined to be 0.66, which implied a possible latent but unmeasured minor construct. Principal component analysis of these root numbers was used to uncover this construct, and the first component PC1 explained 42.6% of the variation in the multivariate root numbers with near equal loadings of all these root numbers. The heritability of this root number-based PC1 was 0.46, greater than the heritability for any single root number phene.

### Root angle phenes for whole crown, main shoot and tillers

Nodal root growth angle of the whole crown ranged from 28 ° to 82 ° from horizontal with an average of 57.5 ° (0 ° is horizontal and 90 ° is vertical). The main shoot had a nodal root growth angle on average of 56.2 ° with tillers 1, 2, and 3 having an average of 55.4 °, 56.6 °, and 55.0 °. The Pearson coefficient was 0.30 between the angle of the main shoot and the angle of tiller 1, while the average inter-item correlation of angles of the whole crown, main shoot, and tillers 1-3 was 0.34. Cronbach’s α was used as a measure of internal consistency of all these angles and was determined to be 0.72, which implied a possible latent but unmeasured major construct. Principal component analysis of these angles was used to uncover this construct, and the first component PC1 explained 47.86% of the variation in the multivariate angles with near equal loadings of all these angles. The heritability of this angle-based PC1 was 0.31, which was again greater than any single shoot root angle phene.

### Relationships among root phenes

Principal component analysis of all phenes of the main shoot and tiller 1 revealed that principal components 1 and 2 explained 24% and 16% of the multivariate variation, respectively, for a total of 40% variation explained (Figure 3). The first component (PC1) was mainly loaded by phenes related to root crown size, such as system width, depth-to-width, and nodal root number. The second component (PC2) was mostly influenced by the angles of the main shoot and tiller 1. The heritabilities of PC1 and PC2 were 0.29 and 0.19, respectively.

Root crown width was found to be positively correlated with shoot number (y = 3.24x + 50.07, R^2^ = 0.13, p <. 001) and total nodal root number (y = 0.64x + 43.04, R^2^ = 0.24, p <.001), and negatively correlated with nodal root growth angle (y = −1.27x + 140.70, R^2^ = 0.33, p <. 001) (Figure 4). The ranges of these four phenes were 28.17 – 196 mm, 2 – 24, 12– 142, and 16.78 – 78.96 ° for root crown width, shoot number, nodal root number, and nodal root angle, respectively. A multiple regression model found that shoot number, total nodal root number, and nodal root growth angle explained 58% of the variation in root crown width (y = −2.6*tiller + −1.22*angle + 1.00*NRN + 113.61, R^2^ = 0.58, p <. 001). Interestingly, in the multiple regression model the slope coefficient for shoot number became negative.

**Fig. 4.**
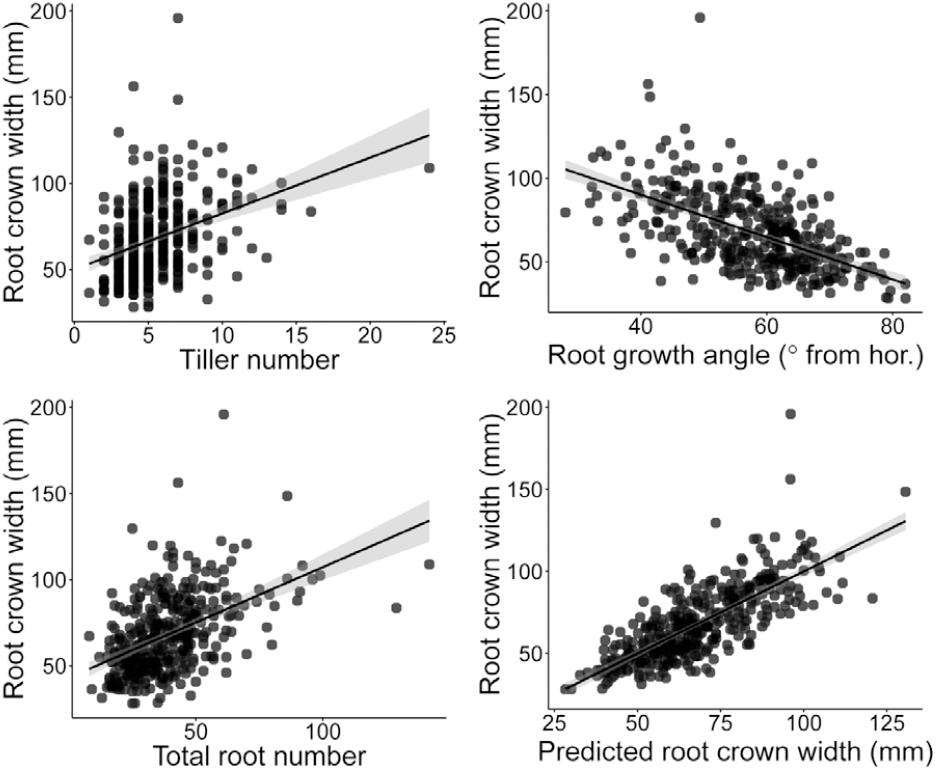
The relationship of root crown width, tiller number, root growth angle, and total root number is revealed in a series of scatter plots. The last panel shows the predicted or fitted values of root crown width from a multiple regression model including the previous three traits as predictors. Every point represents an individual root crown sample. Black line represents the fitted linear regression model and the grey band is the 95% confidence interval.

Nodal root growth angles among the whole crown, main shoot, and tillers were found to be inter-correlated, and the exact relationships were determined through linear regression models (Figure 5). The whole root crown angle was significantly predicted by the main shoot angle (y = 0.53x + 27.49, R^2^ = 0.24, p <. 001), tiller 1 (y = 0.41x + 34.62, R^2^ = 0.16, p <.001), tiller 2 (y = 0.28x + 42.04, R^2^ = 0.09, p <. 001), and tiller 3 (y = 0.32x + 39.92, R^2^ =0.15, p <. 001). A multiple regression model predicting whole root crown angle estimated slopes of the main shoot, tiller 1, tiller 2, and tiller 3 to be 0.35, 0.25, 0.11, and 0.15, respectively, with an intercept of 9.37 (R^2^ = 0.36, p <. 001).

**Fig. 5.**
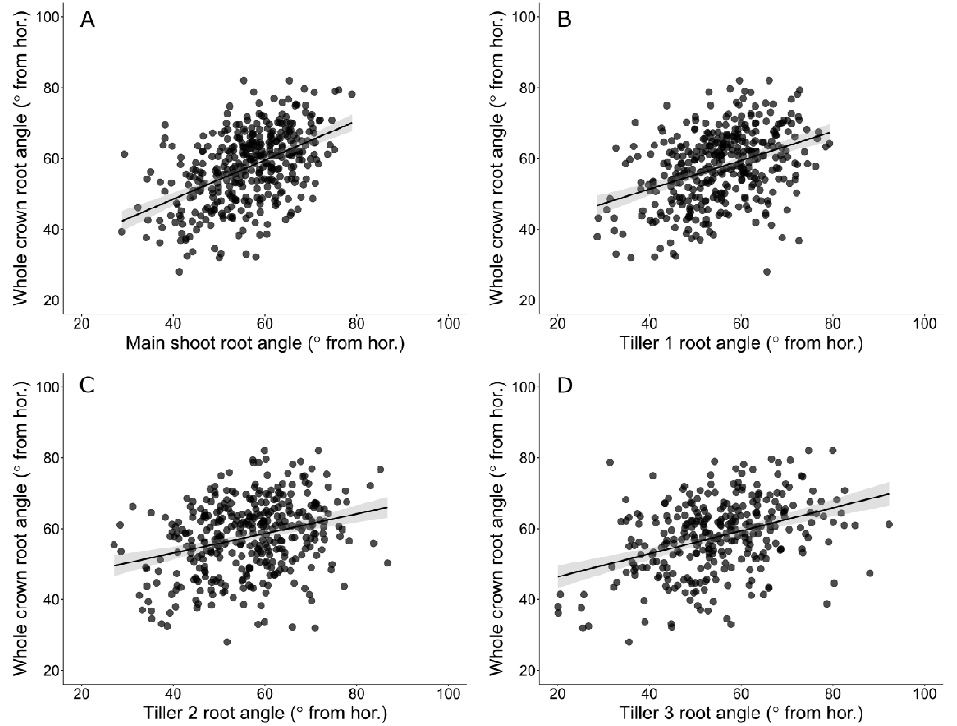
Linear regressions of whole crown root angle versus the root angles of the main shoot (a), tiller 1 (b), tiller 2 (c), and tiller 3 (d). Every point represents an individual root crown sample. Black line represents the fitted linear regression model and the grey band is the 95% confidence interval.

## Discussion

This study applied an intensive phenotyping strategy to excavated root crowns of wheat from the whole crown, and then the individual root crowns separately imaged for the main shoot and tillers. From each of these images, nodal root number, angles, crown root length, and root diameters were extracted. Substantial genotypic variation existed for all measured phenes. Heritabilities were generally greater for whole crown phenes than phenes of individual shoots, with the two greatest being total nodal root number and shoot number at 0.51 and 0.52, respectively. The heritability of root growth angle derived from the whole crown was 0.37. In general, differences in heritabilities for the various phenes may imply they respond more or less to environmental variation. In general, correlations were observed among the same phenes measured across the whole crown, main shoot, and tillers, with inter-item correlation of 0.34 for nodal root growth angle. Principal component analysis of main shoot and tiller 1 phenes revealed a dominant component explaining 24% of multivariate variation and a heritability of 0.29, which was mostly influenced by root crown size-related phenes.

The heritabilities in this study are consistent with others (but often lower) for root system architecture in the literature for wheat ranging between 0.62 and 0.93 (Maccaferri *et al.*, 2016), and between 0.45 and 0.81 in maize (Colombi *et al.*, 2015). Potentially, heritabilities were lower in this study due to only one root crown being sampled per plot rather than several, which may lead to greater variability. Therefore, we suggest sampling three or more root crowns per plot to better estimate the plot level average. Thus, such root crown properties are suitable for breeding programs, although the relation of these phenes to crop productivity are not always known (Lynch, 1995). Linking variation in root phenes to variation in measures of plant performance such as shoot biomass, nutrient content, and yield is crucial for leveraging root phenes in breeding programs. Given the substantial heritabilities, selection of total root number and shoot number could be prioritized.

The relation of root system architecture among shoots in tillering species has not been well-studied. However, understanding this relation would have importance for our understanding root system function. This study demonstrated average inter-item correlations of 0.28 and 0.34 for nodal root number and nodal root growth angle among whole crown, main shoot, and tillers 1-3. Cronbach’s α was found to be 0.66 and 0.72 for numbers and angles among shoots, respectively. In general, these results imply an underlying shared genetic basis among shoots of a plant that causes similar phene states, or as described by Cronbach’sα, an underlying latent construct. Principal components constructed for each phene across shoots had substantial heritabilities, and may be viewed as representations of those latent constructs as revealed by multivariate analysis. Indeed, the heritabilities for the number and angle latent constructs were 0.46 and 0.31, respectively, which were greater for both phenes than measures from any single shoot. Future research may determine whether having different RSA on tillers may have benefits and to what degree the RSA on main shoot and tillers can be decoupled.

Similarly, principal component analysis across all phenes identified components 1 and 2 as representing 24% and 16% of the multivariate variation, respectively. Principal component 1 was predominantly loaded by size-related phenes such as system width, depth-to-width, and nodal root number, while PC2 was mostly influenced by the angles of the main shoot and tiller 1. A similar principal component construction was found for maize root crowns (York and Lynch, 2015), suggesting these relations may be common across species. The heritabilities for PC1 and PC2 were 0.29 and 0.19, respectively. Again, all this suggests not only correlations among phenes, but an underlying genetic basis that causes those correlations, such as a general root vigour driver for PC1. Phenotypic integration has been explored in ecology literature as correlations among phenes due to common developmental pathways, linkage or pleiotropy, and trade-offs in allocation (Murren, 2002). Few studies consider these multivariate relations and underlying genetic drivers, yet they may have great importance for root system function considering the complex integration of root phenes (York *et al.*, 2013).

Root plate spread has previously been hypothesized as important for lodging in wheat (Berry *et al.*, 2000), and was defined as the width of an excavated root system at which the rhizosheath terminated. The root system width reported here is similar. The current study and the Berry *et al*. (2000) study found a strong correlation of root system width with the number of shoots, which limits the usefulness of plate spread for lodging resistance because increased lodging risk is associated with greater numbers of tillers. However, the current research demonstrates that shallow nodal root angles are also greatly positively associated with root system width. Another recent study demonstrated a substantial positive correlation between root plate spread and anchorage strength in the field (Piñera-Chavez *et al.*, 2016). Nodal root number was also positively correlated with root system width in this study, and had substantial heritability. Therefore, nodal root angle and nodal root number deserve further attention as possible sources of increased lodging resistance that are independent of shoot number in wheat.

Several plant phenes in wheat may influence rooting depth. A more vertical angle of seminal roots of wheat seedlings has been linked with more roots at depth in wheat in Australia (Manschadi *et al.*, 2010; Manschadi *et al.*, 2008; Olivares-Villegas *et al.*, 2007). Previous studies in maize also found steeper root angle related to increased rooting depth under low nitrogen field environments in the USA and South Africa (Trachsel *et al.*, 2013). Tillering influences carbon partitioning, and there is some evidence that reduced tillering increases rooting depth in wheat (Duggan *et al.*, 2005; Richards, 2006) and rice (Yoshida and Hasegawa, 1982). Therefore, a plant ideotype with fewer tillers and steeper root angles may be associated with deeper roots.

Although shallow nodal root angles positively correlated with root system width, the effects of number of tillers was as strong. Methods using a basket (Uga *et al.*, 2011) to determine the placement of roots are likely aggregating both phenes which makes resolving genetic and functional associations more difficult (see York *et al.*, 2013 for a discussion of phene aggregates). More direct measurements of angle and other root phenes as conducted here are likely to benefit plant breeding.

The relation of root system architecture to crop performance in wheat is partially established, but many gaps in knowledge remain. A recent study found a positive correlation of root number with enhanced growth in compacted soil (Colombi and Walter, 2017). QTLs for root angle and number overlapped with QTLs for yield in a wheat mapping population (Canè *et al.*, 2014), and again in another study (Maccaferri *et al.*, 2016). Recently, a major vernalization gene was also shown to influence seminal root growth angle and possibly nodal root growth angle in wheat and barley, providing an intriguing example of shared control of shoot and root properties (Voss-Fels *et al.*, 2017).

Excavating in the field took 2 days for a team of three, while washing the root crowns took 3 days for a team of four. Imaging took 5 days with two users, and the manual image analysis took 3 days for a single user. Automation of all steps would benefit throughput, which would allow more replicates and greater population sizes that ultimately allow greater statistical power in resolving genetic regions and functional relations of root phenes to crop performance. Root crown image analysis tools like REST (Colombi *et al.*, 2015) are part of the solution, however no root crown tool counts axial roots even though some extracted features correlate to numbers. Furthermore, acquiring root crown images under reproducible lighting conditions yielding high contrast images of roots against background has been challenging and leads to some failures of segmentation and analysis. More custom hardware solutions for root crown imaging would be useful for the community. Partial automation of excavation of root crowns and removal from the field, along with automated soil removal, would also increase throughput. Root crown phenotyping is ripe for technological advancement that will undoubtedly contribute to the understanding of root system function.

Intensive phenotyping of root crowns in wheat has revealed correlations among shoots of the same plant for root crown phenes and substantial heritability of root crown phenes. Multivariate methods identified unmeasured latent constructs that may drive variation in several phenes simultaneously, perhaps through pleiotropic effects of interacting genes. Applying these methods in breeding populations and simultaneously measuring crop performance will allow inferring how root phenes contribute to yield, but requires advancements in throughput. Combining optimized RSA with other properties that influence aspects of the holistic rhizosphere (York *et al.*, 2016) will likely have synergistic effects due to phene integration (York *et al.*, 2013). Recently, this method was used to demonstrate a positive influence of nodal root number and growth angle on both root depth and yield of wheat in the field (Slack *et al.*, 2018). Root crown phenotyping in wheat has substantial potential for contributing to the global breeding effort for increased agricultural sustainability and food security.

## Acknowledgments

This work was funded by FP7-IDEAS-ERC FUTUREROOTS: Redesigning root architecture for improved crop performance ERC Future Roots Project ID: 294729. We thank Limagrain UK Ltd and John Innes Centre, UK for the use of the Savannah × Rialto DH population material.

